# Fast and Accurate Influenza Forecasting in the United States with Inferno

**DOI:** 10.1101/2021.01.06.425546

**Authors:** Dave Osthus

## Abstract

Infectious disease forecasting is an emerging field and has the potential to improve public health through anticipatory resource allocation, situational awareness, and mitigation planning. By way of exploring and operationalizing disease forecasting, the U.S. Centers for Disease Control and Prevention (CDC) has hosted FluSight since the 2013/14 flu season, an annual flu forecasting challenge. Since FluSight’s onset, forecasters have developed and improved forecasting models in an effort to provide more timely, reliable, and accurate information about the likely progression of the outbreak. While improving the predictive performance of these forecasting models is often the primary objective, it is also important for a forecasting model to run quickly, facilitating further model development, improvement, and scalability. In this vein I introduce Inferno, a fast and accurate flu forecasting model inspired by Dante, the top performing model in the 2018/19 FluSight challenge. When compared to all models that participated in FluSight 2018/19, Inferno would have placed 2nd in both the national and state challenges, behind only Dante. Inferno, however, runs in minutes and is trivially parallelizable, while Dante takes hours to run, representing a significant operational improvement with minimal impact to performance. A future consideration for forecasting competitions like FluSight will be how to encourage improvements to secondarily important properties of forecasting models, such as runtime, generalizability, and interpretability.

## 1 Introduction

Infectious disease outbreaks can be disruptive, deadly, and complex. By the end of November 2020, COVID-19 had killed almost 1.5 million people globally and over 250 thousand people in the United States (U.S.) [6]. Each year in the U.S., seasonal influenza kills tens of thousands of people and hospitalizes hundreds of thousands [25]. Life saving resources, such as respirators, antivirals, vaccines, and medical professionals must be allocated to ensure locations are prepared and ready for the impending outbreak.

This is where infectious disease forecasting comes in. If forecasts can reliably anticipate the progression of an outbreak, we may be better prepared to confront it when it arrives. Infectious disease forecasting is still relatively young, but can no longer claim novelty. There has been a flurry of infectious disease forecasting challenges/collaborations in the last ten years, including the Defense Advanced Research Projects Agency’s 2014/15 Chikungunya challenge [5], a collection of vector-borne disease challenges hosted by the U.S. Centers for Disease Control and Prevention (CDC) for dengue (2015) [9], Aedes (2019) [23], and West Nile virus (2020) [24], the U.S. CDC COVID-19 forecasting collaboration (2020) [27], and the U.S. CDC’s flagship influenza forecasting challenge, FluSight, held annually since the 2013/14 flu season. The FluSight challenge alone has resulted in a wave of infectious disease forecasting model development [13, 12, 11, 14, 20, 19, 4, 3, 10, 15, 1, 28, 26].

The organizing body of a forecasting challenge (in the case of FluSight, the U.S. CDC) provides immense operational and research value by determining forecasting targets of public health relevance through interactions with their state and local public health partners, identifying relevant data sources and making them publicly available to forecasters, and defining the forecast evaluation criteria — a more challenging task than it may first appear (see [2] and [21]).

For instance, the FluSight challenge asks forecasters to predict seven targets on a weekly basis throughout the flu season: 1 through 4-week-ahead forecasts of influenza-like illness (ILI), the week of flu season onset, the week the flu season will peak, and the peak value of ILI for the flu season. Forecasts are made for states, Health and Human Services (HHS) regions, and the United States. ILI data collected by the U.S. Outpatient Influenza-like Illness Surveillance Network (ILINet) are used for forecasting; targets are defined as summaries of ILI data. FluSight uses the log scoring rule to evaluate forecasts. The log scoring rule evaluates probabilistic forecasts, requiring forecasters to not only provide a prediction of what they think will happen in the future but also quantify how sure they are of that. The choice made by the U.S. CDC to use a log scoring rule makes clear their position that uncertainty quantification is of value to public health. Given a set of forecasting targets and an evaluation metric, forecasters develop models capable of forecasting the targets with the goal of maximizing their forecast evaluation score.

While a forecasting model’s predictive performance is and should be of primary importance, it is not exclusively important. This seems obvious upon even cursory consideration. For instance, all else equal, an interpretable forecasting model is better than a black box fore-casting model. All else equal, a generalizable forecasting model applicable to many disease forecasting contexts is better than a highly-tailored model to a specific disease context. All else equal, a forecasting model that runs quickly and is scalable is better than one that is slow and computationally expensive. While all of these seem obvious, none of these secondary factors are incorporated into FluSight’s forecast evaluation criteria; it *only* measures the predictive performance of the model. As a result, much of the forecasting research of the past decade has focused more on developing models that improve forecasting scores and less on developing models that are generalizable, interpretable, scalable, and fast.

In this paper, I focus on improving the runtime of flu forecasting models while maintaining high prediction standards with the presentation of Inferno, a fast and accurate flu forecasting model. Inferno is a parallelizable, empirical Bayesian forecasting model inspired by Dante, the top performing model in FluSight 2018/19 [14]. The achieved goal of Inferno is to maintain the high predictive performance of Dante but substantially decrease the run-time. As will be discussed later, Inferno would have placed 2nd only to Dante in the 2018/19 FluSight challenge, but runs in minutes rather than hours, constituting a significant speed-up in operational performance.

In the remainder of this paper, I describe the details to Inferno (Section 2) and present Inferno’s forecasting performance as compared to all participating models in FluSight 2018/19 (Section 3). I conclude the paper by raising important questions the infectious disease forecasting community must grapple with in order to improve the utility for forecasting challenges for public health.

## 2 Methods

### 2.1 Dante Background

Dante is a multiscale, probabilistic, influenza forecasting model. Dante has two sub-models: a state forecasting model and an aggregation model which combines state forecasts to produce HHS regional and United States forecasts. The state forecasting model is a statistical model where the expectation of ILI on a given week, state, and season is modeled as a function of four components: an overall trend component, a state-specific deviation component, a season-specific deviation component, and a state/season-specific deviation component. These four components are each modeled as random or reverse-random walks — flexible time series models that capture temporal correlation. By modeling all states and past flu seasons jointly, Dante achieves self-consistency and is able to borrow information across seasons and space. By modeling the HHS regional and United States forecasts as U.S. Census weighted averages of state forecasts, Dante ensures self-consistency across geographic scales. For more details on Dante, see [14].

Dante is a fully Bayesian model, capturing uncertainty in all model parameters, latent states, and forecasts through its posterior (predictive) distribution. The fully Bayesian formulation and self-consistency of Dante comes at a computational price, however. Dante represents a large model that will grow each year as more historical data are added and is not well-positioned to scale with possible future changes/expansions to FluSight (e.g., county-level forecasting). Nothing is precomputed and due to its interconnected model structure, it is not obvious how to break up Dante to exploit parallelization.

Inferno was developed to addresses these computational shortcomings. Inferno is an empirical Bayesian analogue to the fully Bayesian Dante, where instead of modeling historical flu seasons directly, Inferno uses historical flu seasons to precompute model parameters. Inferno trades in self-consistency for parallelization, allowing all states, HHS regions, and the United States to be fit independently. In Section 2.2, I describe the Inferno forecasting model.

### 2.2 Inferno

Let *y*_*t*_ ∈ (0, 1) for *t* = 1, 2, …, *T* be ILI*/*100 for states or state-weighted ILI*/*100 for HHS regions or the United States (collectively referred to as (w)ILI) for week of season *t*, where *t* = 1 corresponds to Morbidity and Mortality Weekly Report (MMWR) week 40, roughly the beginning of October, and *T* = 35 roughly corresponds to the end of May. Inferno’s generative model is defined as follows:

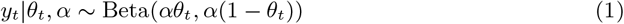

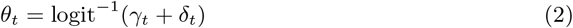

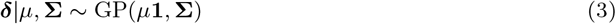

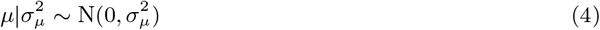

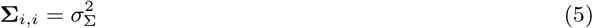

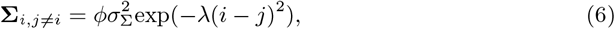

where *y*_*t*_ is the noisy but observable measurement of ILI*/*100 on week *t, θ*_*t*_ is the true but unobservable value of ILI*/*100 on week *t*, ***δ*** = (*δ*_1_, *δ*_2_, …, *δ*_*T*_)^*′*^ is a *T* × 1 vector, **1** is a *T* × 1 vector of 1s, **Σ** is a *T* × *T* positive semi-definite matrix, GP(***µ*, Σ**) is a Gaussian process (GP) with mean ***µ*** and covariance **Σ**, the scalar parameters *α*, 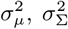,*λ* are all greater than 0, and *ϕ* ∈ [0, 1]. In this paper, bold quantities represent vectors or matrices, while non-bold quantities represent scalars. The Beta distribution of Equation 1 requires *y*_*t*_ ∈ (0, 1). Thus, all *y*_*t*_ below a low threshold *l* are set equal to *l* and all *y*_*t*_ above 1 − *l* are set to 1 − *l*. For this work, *l* = 0.0005.

Inferno takes an empirical Bayesian approach, where unknown parameters are estimated from historical training data. The following outlines a six step procedure to estimate the unknown parameters *α*, ***γ*** = (*γ*_1_, *γ*_2_, …, *γ*_*T*_)^*′*^, 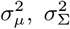, *λ*, and *ϕ* and to use Markov chain Monte Carlo (MCMC) to sample and forecast from Inferno’s posterior predictive distribution.

#### 2.2.1 Step 1: Estimate *θ*_*s,t*_

For a given geographical unit (e.g., state, region country), let *y*_*s,t*_ by (w)ILI for training season *s* ∈ 1, 2, …, *S* and week of season *t*. Fit 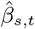 as a 3-week moving average:

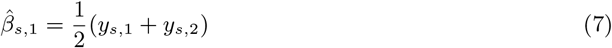

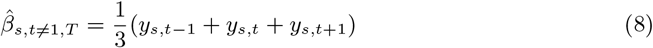

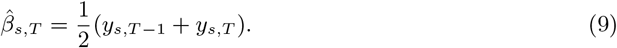

Figure 1 shows the moving average fit to ILI in Illinois. By construction, the moving average captures the shape of the ILI curve. The moving average, however, can miss sharp changes in ILI caused by differences in reporting practices over holidays. For instance, we see that the moving average most often underestimates ILI the week of Christmas (*t* = 13, or MMWR week 52).

**Figure 1:**
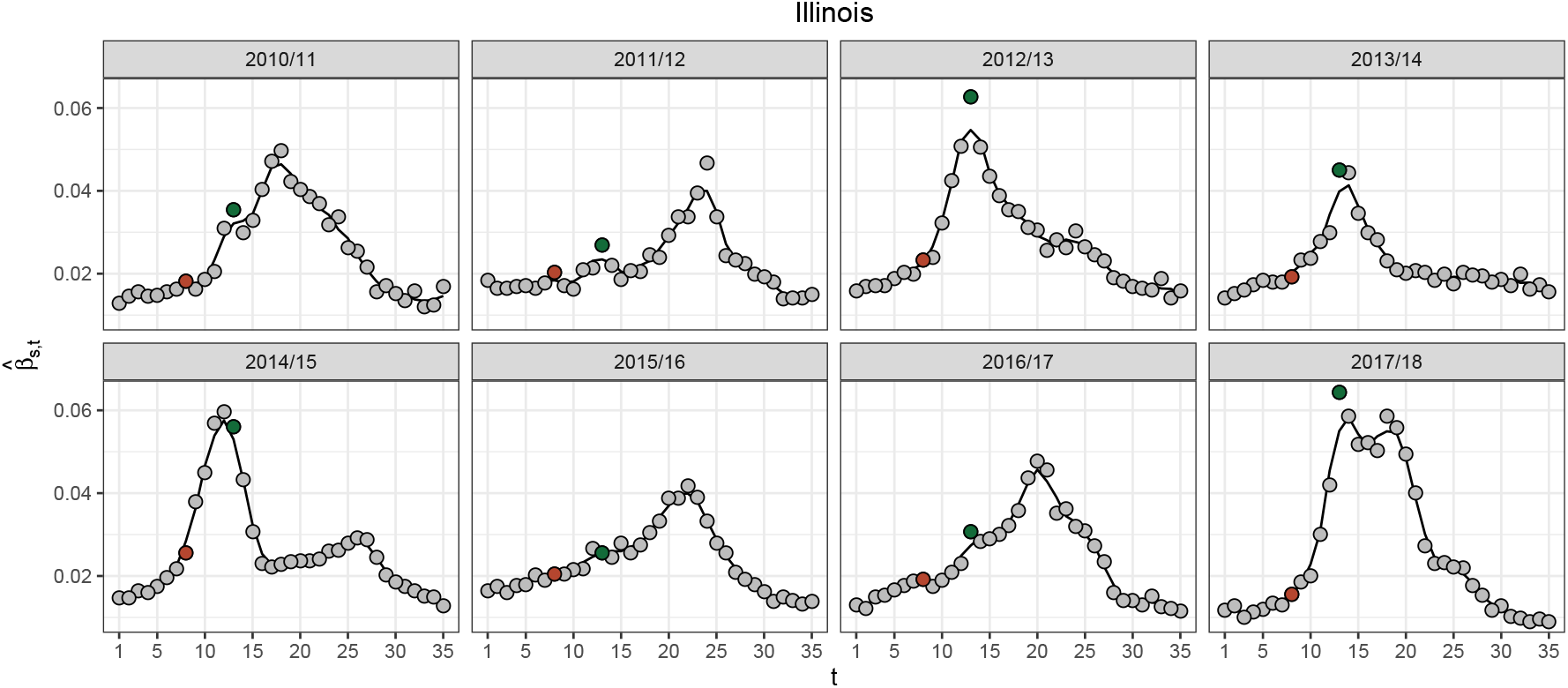
ILI (grey points) and 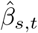 (black line) for the historical seasons for Illinois. ILI for the week of Thanksgiving (*t* = 8) and Christmas (*t* = 13) are highlighted in brown and green, respectively. 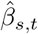 typically underestimates the sharp uptick in ILI observed on Christmas and to a lesser extent Thanksgiving, which is likely a result of changes in reporting and care-seeking behavior over the holidays.

To capture the systematic sharp changes in ILI that are common across training seasons, Inferno computes the quantity *τ*_*t*_:

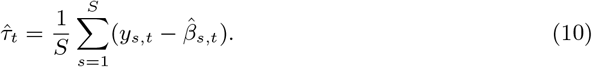

Figure 2 plots *τ*_*t*_ for all states. *τ*_*t*_ captures the holiday effects in ILI, with a small but consistent positive *τ*_*t*_ on the week of Thanksgiving (*t* = 8, or MMWR week 47) and a larger positive effect the week of Christmas.

**Figure 2:**
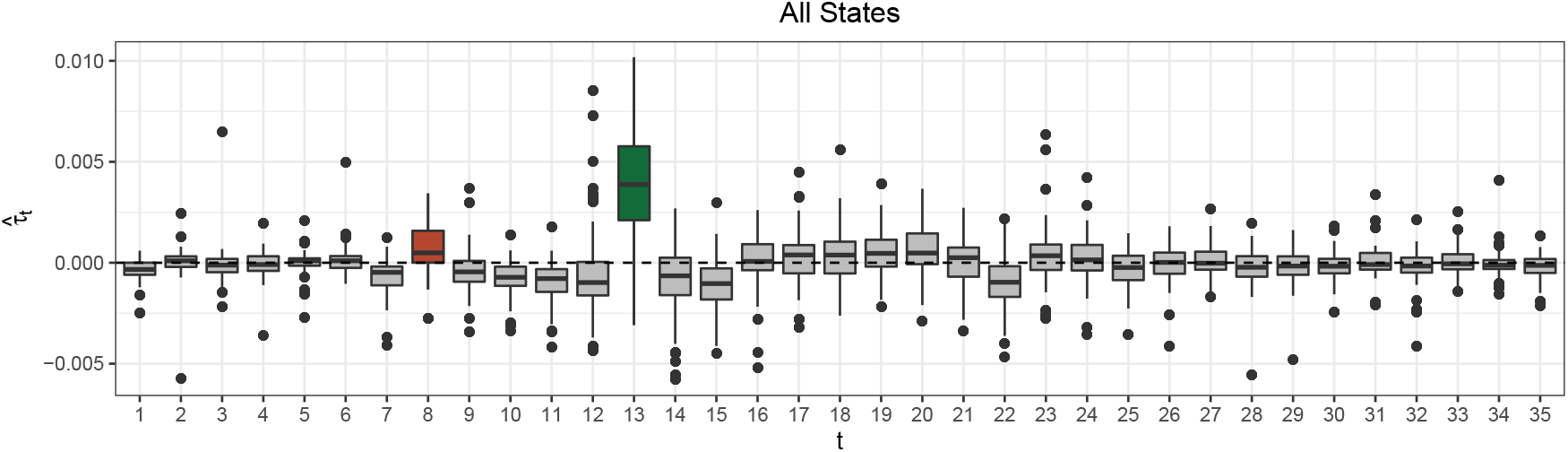
The quantity 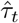 for all states. 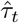 the week of Thanksgiving (brown) and Christmas (green) are systematically positive, likely as a result of systematic changes to reporting and care-seeking behavior over the holidays.

Finally, the quantity *θ*_*s,t*_ captures both the ILI profile 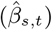 and the holiday effects 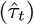:

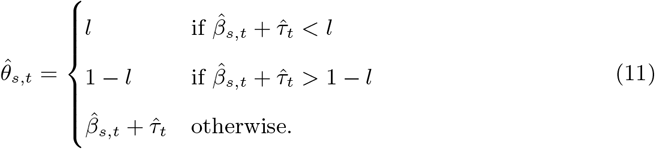

Figure 3 shows how 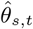 tracks the profile of the ILI season, like 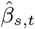, but better tracks ILI on the holidays, especially Christmas.

**Figure 3:**
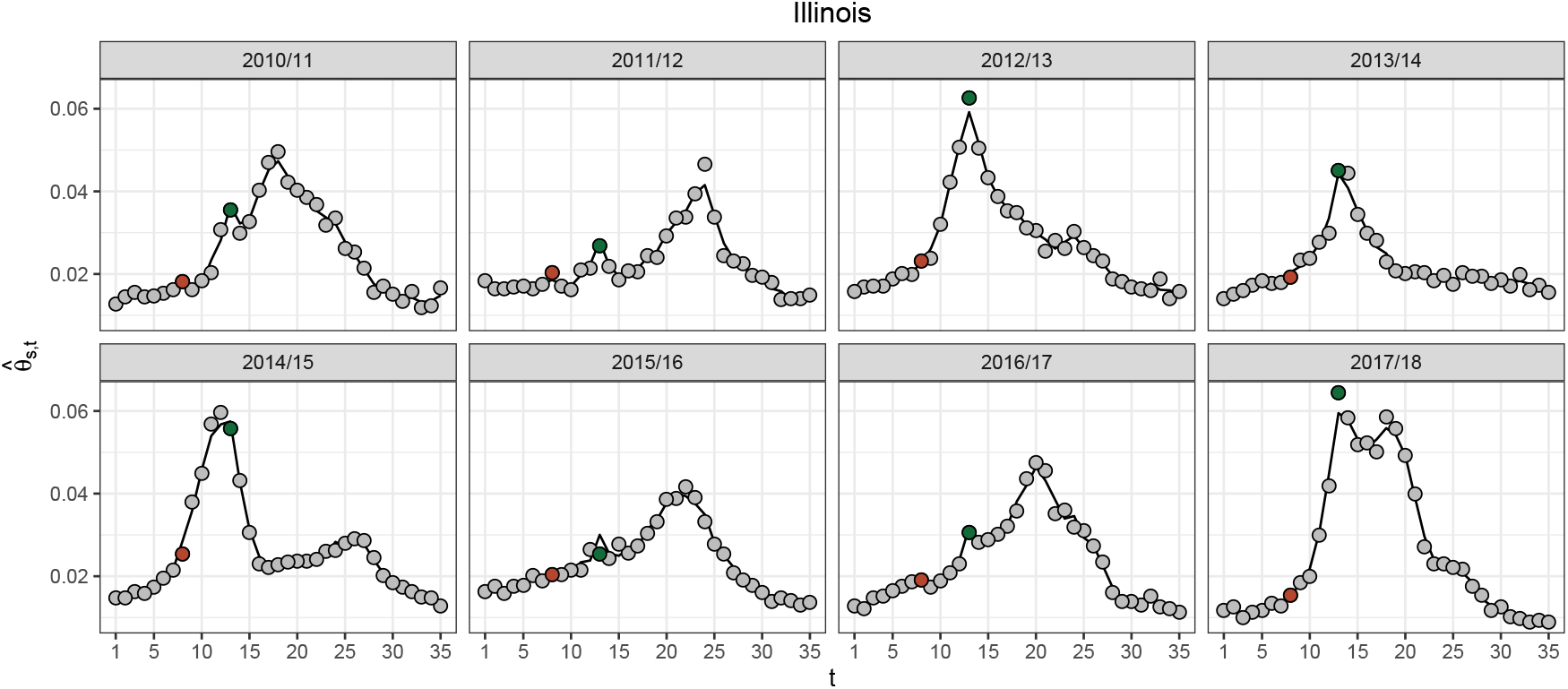
ILI (grey points) and 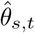 (black line) for the historical seasons for Illinois. ILI for the week of Thanksgiving (*t* = 8) and Christmas (*t* = 13) are highlighted in brown and green, respectively. 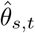 better matches ILI data on the holidays than 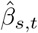 (Figure 1) by accounting for the systematic reporting and care-seeking changes over the holidays, as accounted for by 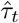.

#### 2.2.2 Step 2: Estimate *α*

Inferno estimates 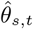 in order to facilitate the estimation of the other unknown quantities of Inferno’s generative model. The expectation and the variance of Inferno’s data model (Equation 1) are,

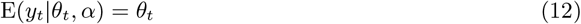

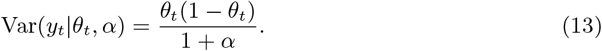

The parameter *α* controls the variance of the data model, capturing the week-to-week variability in the ILI data. The larger *α* is, the smaller the variance reflecting less week-to-week noise in the ILI data. The smaller *α* is, the larger the variance reflecting more week-to-week noise in the ILI data. We estimate *α >* 0 as the maximum likelihood estimate (MLE) of Inferno’s data model by minimizing the negative log likelihood:

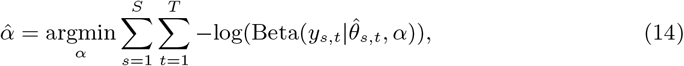

where log(*x*) is the natural log of *x*,

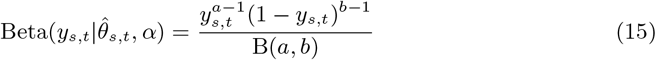

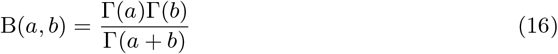

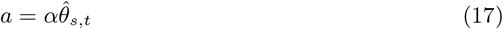

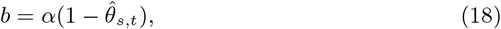

and Γ() is the gamma function.

The top of Figure 4 shows 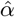 for all states, territories, and cities (collectively referred to as states). States like the U.S. Virgin Islands, North Dakota, and Puerto Rico have the smallest 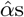, reflecting they have the largest week-to-week noise in their ILI data, while states like California, Illinois, and New York City have the largest 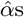, reflecting they have the smallest week-to-week noise in their ILI data. The bottom of Figure 4 shows the 95% prediction interval for Beta 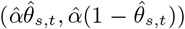 for North Dakota, Nevada, and Illinois, illustrating the different levels of week-to-week noise in ILI data across states.

**Figure 4:**
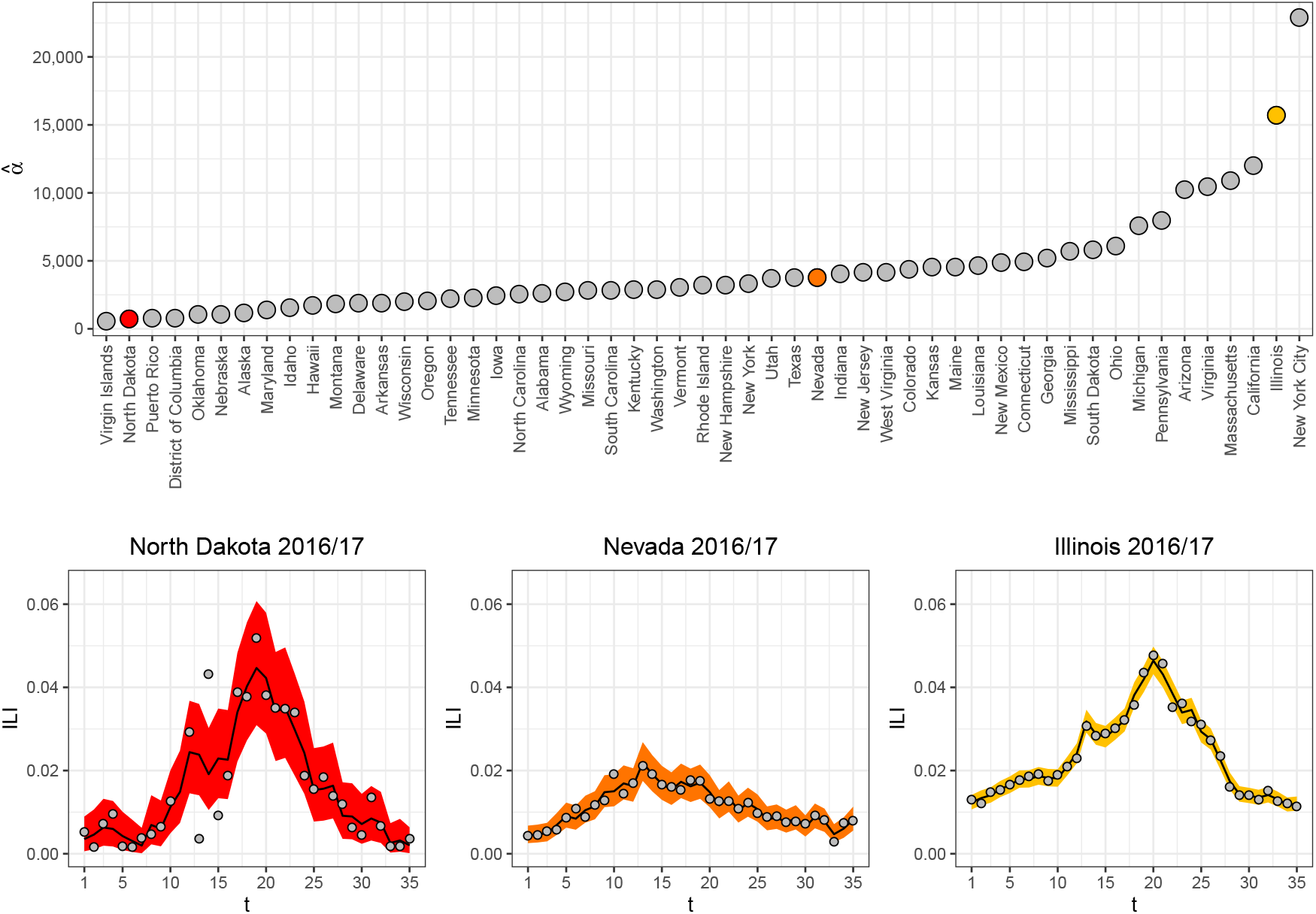
(Top) 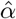 for all states based on training data from 2010/2011 through 2017/18. (Bottom) ILI (grey points), 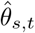 (black line) and 95% prediction interval for Beta 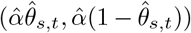 (ribbon) for North Dakota, Nevada, and Illinois in 2016/17. 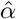 captures the week-to-week noise in ILI data that systematically varies from state-to-state, where North Dakota has more week-to-week noise than Illinois.

#### 2.2.3. Step 3: Estimate *γ*_*t*_

Seasonal flu has a typical shape to it in the United States. ILI starts at low levels early in the season, rises to a peak between December and March, and reverts to low levels by the end of May. The role of ***γ*** is to capture this typical seasonal flu profile. Inferno computes *γ*_*t*_ as follows:

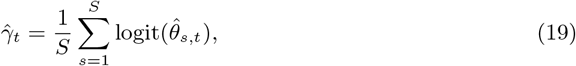

where logit(*p*) = log(*p/*(1 − *p*)).

Figure 5 shows 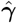 for North Dakota, Nevada, and Illinois. We see for all states, 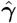 captures the typical profile of seasonal flu on the logit scale, with low levels at the beginning of the flu season, ramping up to a peak in the middle, then reverting back to low levels by the end.

**Figure 5:**
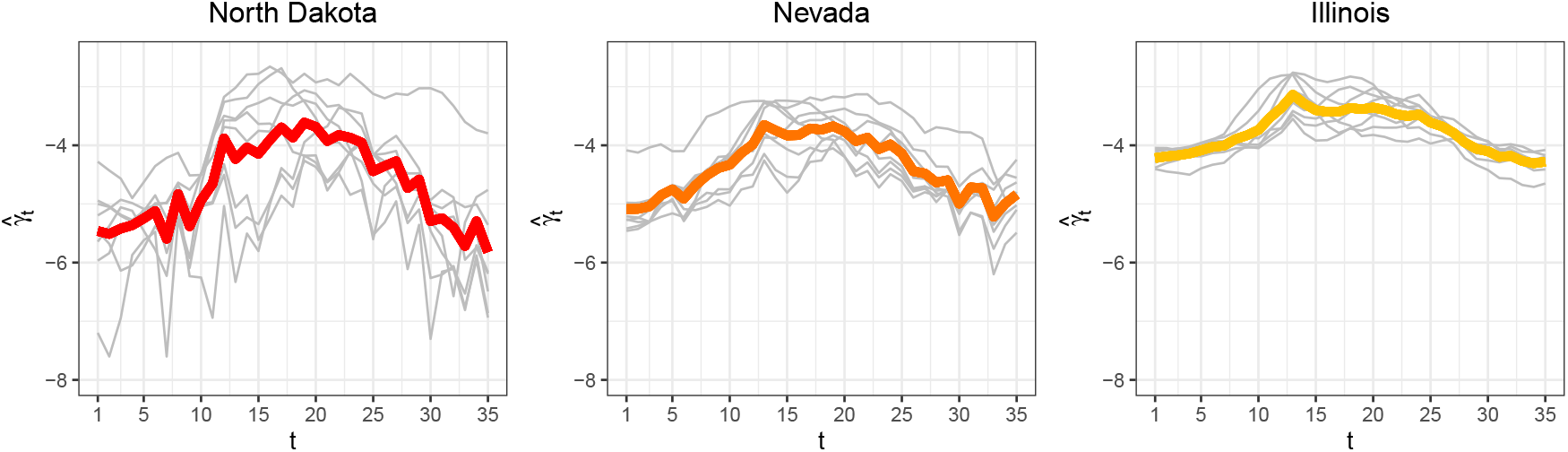
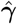 (colored line) and 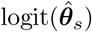 (grey lines) for North Dakota, Nevada, and Illinois. 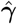 captures the typical profile of seasonal flu specific to each state on the logit scale.

#### 2.2.4 Step 4: Estimate 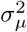

Equation 2 is the mean of Inferno’s data model. While ***γ*** captures the typical profile of seasonal flu, ***δ*** captures season-specific deviations from ***γ***. Inferno models ***δ*** with a Gaussian process (GP), a stochastic process where any finite collection of random variables has a multivariate normal distribution. That is,

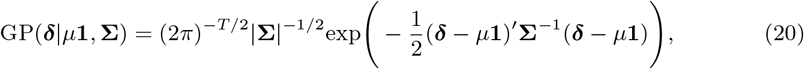

where **1** is a *T* × 1 vector of ones, **Σ** is a *T* × *T* positive semi-definite matrix, |**Σ**| is the determinant of **Σ**, and **Σ**^*−*1^ is the inverse of **Σ**. The model for the mean of the GP *µ* is

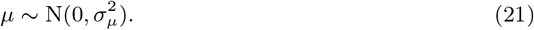

Step 4 describes how to estimate 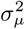.

First compute the following quantities:

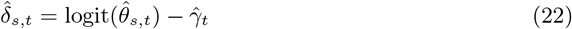

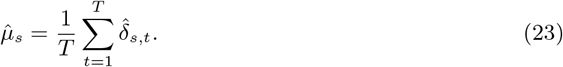

The top of Figure 6 shows 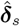 and 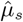 for North Dakota, Nevada and Illinois. The quantity 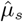 captures how far, on average, 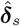 deviates from **0**.

**Figure 6:**
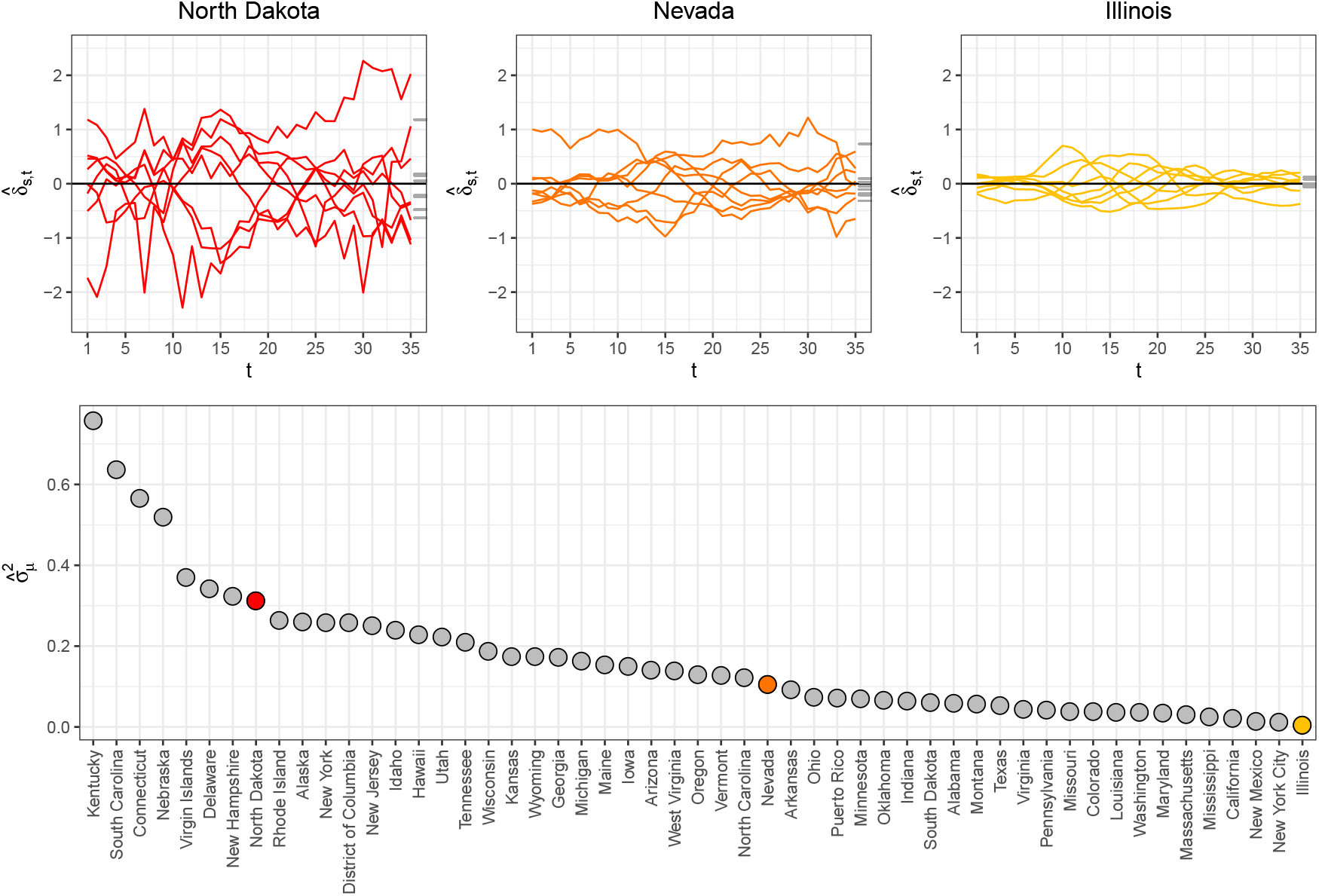
(Top) 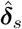 (colored lines) and 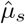 (grey tick marks) for North Dakota, Nevada, and Illinois. North Dakota exhibits more season-to-season variability in 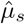 than Illinois, as can be seen in the spread of 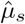. (Bottom) 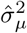 for all states. Considerable variation in 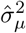across states is observed.

The quantity 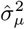 is computed as the unbiased sample variance of 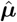:

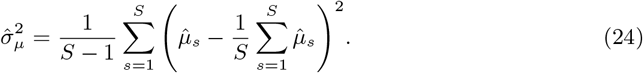

The bottom of Figure 6 shows 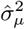 for all states. Some states, like North Dakota, have appreciable average season-to-season variation while other states, like Illinois, have smaller average season-to-season deviations from their typical seasonal flu profiles.

#### 2.2.5 Step 5: Estimate 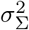, *λ, ϕ*

Step 5 estimates the covariance parameters in **Σ**. The covariance matrix captures different characteristics of ***δ***. Recall Equations 5 and 6:

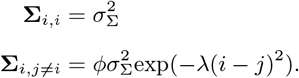

The parameter 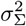 is the marginal variance for the GP. It captures how far ***δ*** − *µ***1** typically deviates from 0. The top of Figure 7 plots 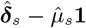 for North Dakota, Nevada, and Illinois. North Dakota exhibits more variability than Illinois as can be seen with its wider range of values. Inferno estimates 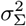 as

**Figure 7:**
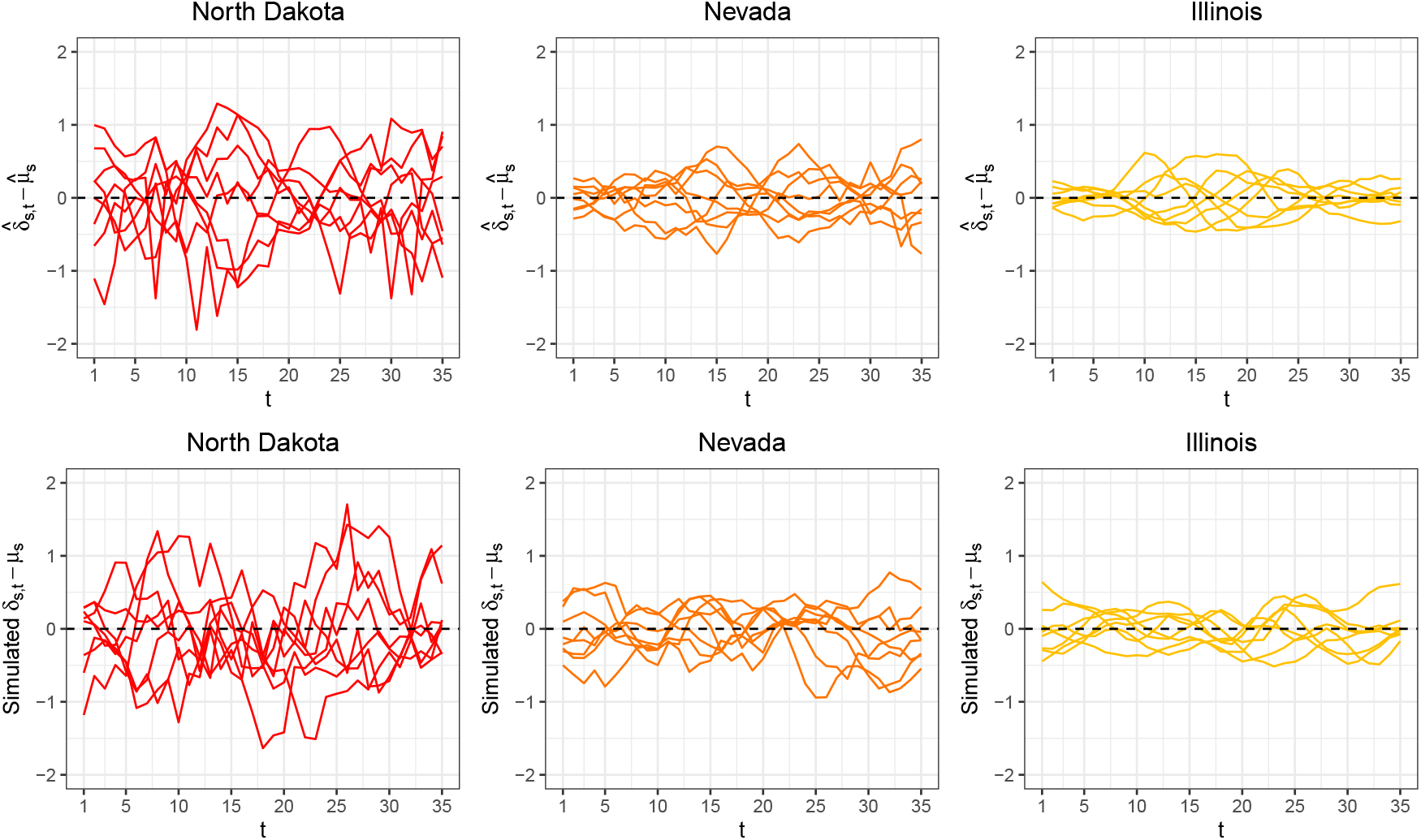
(Top row) The estimated times series from training data for the quantities 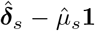. (Bottom row) Realizations drawn from GP(**0**, 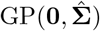). Good visual agreement is seen between the simulated ***δ***_*s*_ *− µ*_*s*_**1** and the 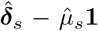 calculated from training data, suggesting the Gaussian process is able to capture heterogenous discrepancy characteristics across states.

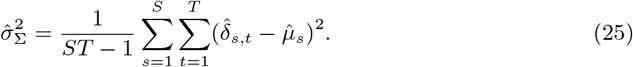

The remaining parameters of **Σ** are *ϕ* and *λ*. They collectively capture two different characteristics of ***δ***. The parameter *ϕ* captures the smoothness of ***δ***. For instance, 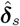 for Illinois in Figure 6 are much smoother than 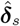 for North Dakota. The parameter *ϕ* captures this feature, with *ϕ* close to 1 resulting in smoother ***δ***s. The second characteristic of ***δ*** captured by *ϕ* and *λ* is the correlation between entries of ***δ***. For instance, the correlation between *δ*_*i*_ and *δ*_*i*+1_ is

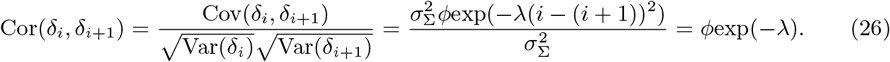

Inferno estimates *ϕ* and *λ* by minimizing the negative log likelihood:

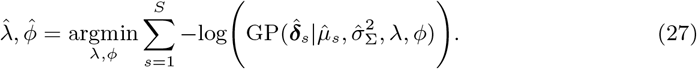

Figure 8 plots the covariance parameter estimates for all states. North Dakota has larger marginal variance (larger 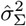), less smoothness (smaller 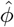), and lower correlation (smaller 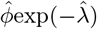) than Illinois.

**Figure 8:**
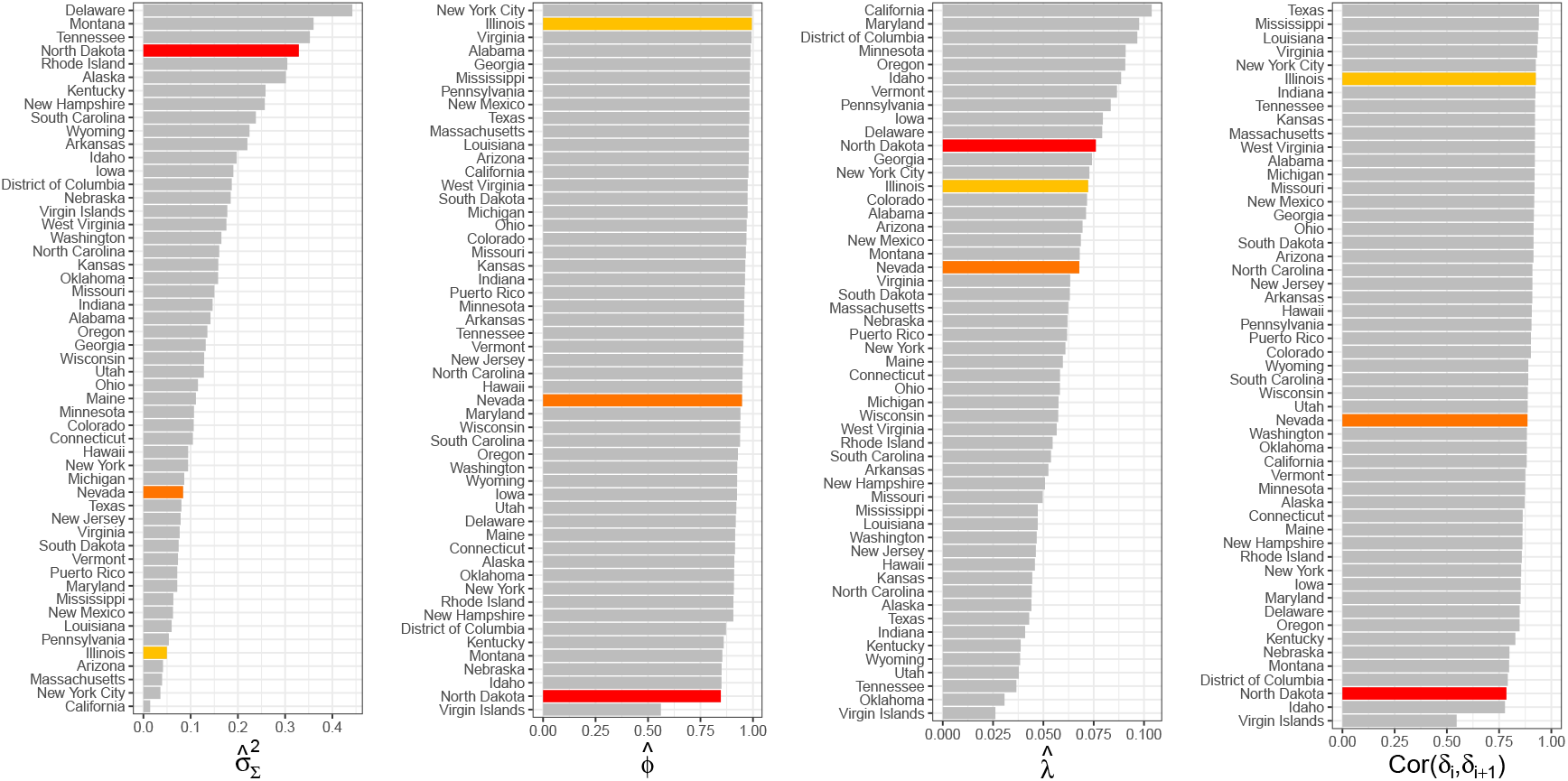
Estimated GP covariance parameter estimates for all states. Parameter estimates for North Dakota, Nevada, and Illinois are highlighted in red, orange, and yellow, respectively. North Dakota has larger marginal variance (larger 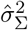), less smoothness (smaller 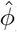), and lower correlation (smaller Cor(*δ*_*i*_, *δ*_*i*+1_)) than Illinois.

The bottom of Figure 7 shows realizations drawn from 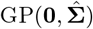. The fitted GP does a good job capturing the different characteristics of the empirical quantities 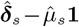, suggesting the GP is a defensible generative model for ***δ***.

#### 2.2.6 Step 6: Sample Forecasts from Inferno

The sixth and final step of Inferno is to replace parameters with their estimates and sample from the posterior predictive distribution. Specifically, the generative model with parameters replaced by their estimates is

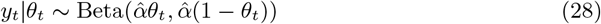

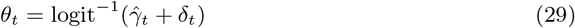

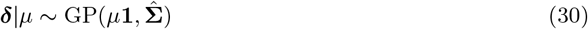

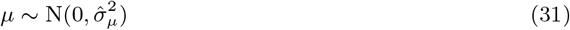

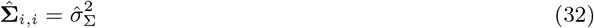

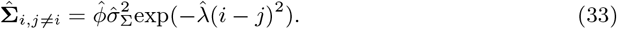

Inferno forecasts the entire flu season by sampling from the posterior predictive distribution given the first *t* weeks of ILI observations:

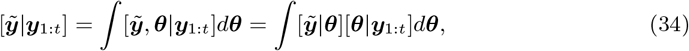

where [*X*|*Y*] is the conditional distribution of *X* given *Y* and 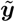 is assumed to be independent of ***y***, given ***θ*** where ***θ*** generically represents all parameters and latent states of Inferno. The posterior predictive distribution of Equation 34 is not known in closed form. Markov chain Monte Carlo (MCMC) sampling is used to draw from the posterior predictive distribution. The probabilistic programming language JAGS (Just Another Gibbs Sampler) [16] is used to execute the MCMC sampling. JAGS is called with functions from the rjags package [17] in the programming language R [18]. The results are *J* draws from the posterior predictive distribution of Equation 34. For this paper, forecasts are based on *J* = 25, 000 draws, discarding the first 12,500 draws as burn-in and thinning the remaining 12,500 draws by two, resulting in forecasts based on 6,250 MCMC draws. The JAGS code that implements Inferno can be found in Appendix A.

Figure 9 shows the forecasts for North Dakota, Nevada, and Illinois throughout the 2018/19 flu season. The posterior predictive mean and the 95% posterior prediction interval are shown as summaries of the forecasts.

**Figure 9:**
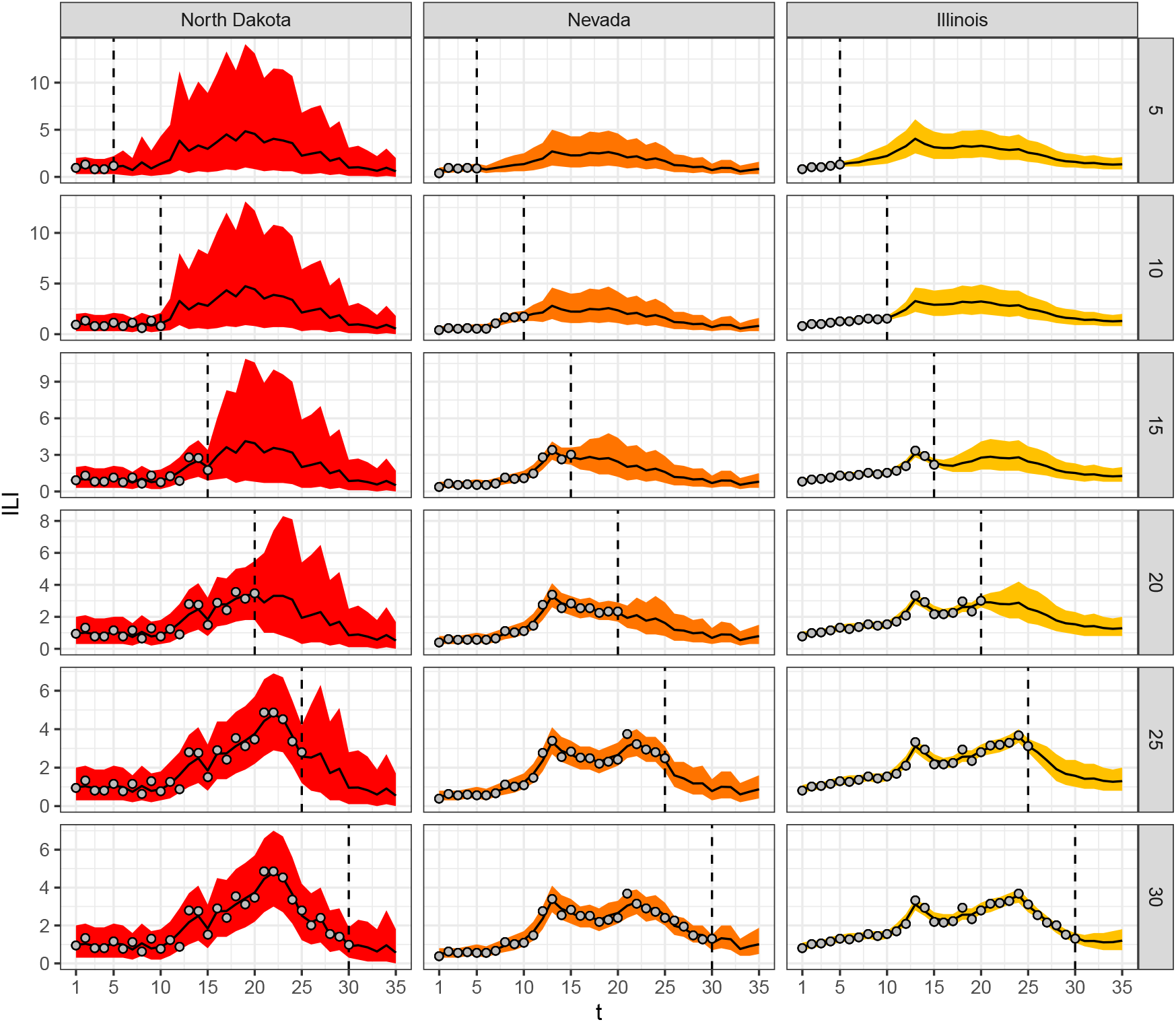
Inferno forecasts for the 2018/19 flu season for North Dakota, Nevada, and Illinois (columns) made *t* = 5, 10, 15, 20, 25, 30 weeks into the flu season based on summaries of draws from the posterior predictive distribution 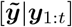 of Equation 34 (rows). Posterior mean (black line) and 95% prediction intervals (ribbons) are displayed, along with ***y***_1:*t*_ (grey points).

## 3 Results

Inferno is compared to all models that participated in the U.S. CDC’s 2018/19 National and Regional FluSight challenge as well as the State challenge. Forecasting follows the guidelines outlined by the CDC FluSight challenge; see [22] for details. The forecasts and the evaluation procedure is briefly described below.

Forecasts are made for four short-term targets (1, 2, 3, and 4-week-ahead) and three seasonal targets (the peak week, the peak percentage, and the onset week — onset is not forecasted for the State challenge). All forecast targets are binned and a probability is assigned to each bin such that the sum of all probabilities over all bins for a target equals 1. Say bin *b* is the bin containing the correct target and *p*_*b*_ ∈ [0, 1] is the probability assigned to the correct bin. The log score is then max(−10, log(*p*_*b*_)). When *b* is the bin of the correct target, the evaluation criteria is called the *single-bin log score*; single-bin log score is the scoring criteria used starting with the 2019/20 FluSight challenge and is a proper score. When *b* is the set of bins containing the correct target plus/minus a set of predefined neighboring bins, the evaluation criteria is called the *multi-bin log score*; multi-bin log score is the scoring criteria used in the 2018/19 FluSight challenge. The multi-bin log score is an improper scoring rule [21]. Multi-bin skill and single-bin skill are derived by exponentiating the multi-bin and single-bin log scores, respectively.

ILI data is subject to weekly revisions. As a result, it is important to use the ILI estimates that were available at the time to make faithful comparisons to models that participated in the real-time FluSight challenges. Data available at historical dates are made available by the Carnegie Mellon University Delphi group’s API [7] and were used to produce the results in this section.

Figure 10 and Table 1 show the multi- and single-bin skills for Inferno and all models that participated in the 2018/19 FluSight challenges. Inferno would have placed 2nd only to Dante in the 2018/19 FluSight National and Regional as well as State challenges. FluSight 2018/19 used multi-bin skill as the forecast evaluation. Starting with FluSight 2019/20, single-bin skill will be used. While single-bin and multi-bin skills are correlated, as can be seen in Figure 10 the relationship is not perfect. Models can rise or fall in the relative ranking depending on which evaluation metric is used for scoring, highlighting that the evaluation metric the forecasting challenge organizing body selects is of consequence. Inferno and Dante both perform better under the multi-bin skill evaluation than single-bin skill, but are both top 4 models by either evaluation metric. Most importantly, the drop in predictive performance from Dante to Inferno is small.

**Figure 10:**
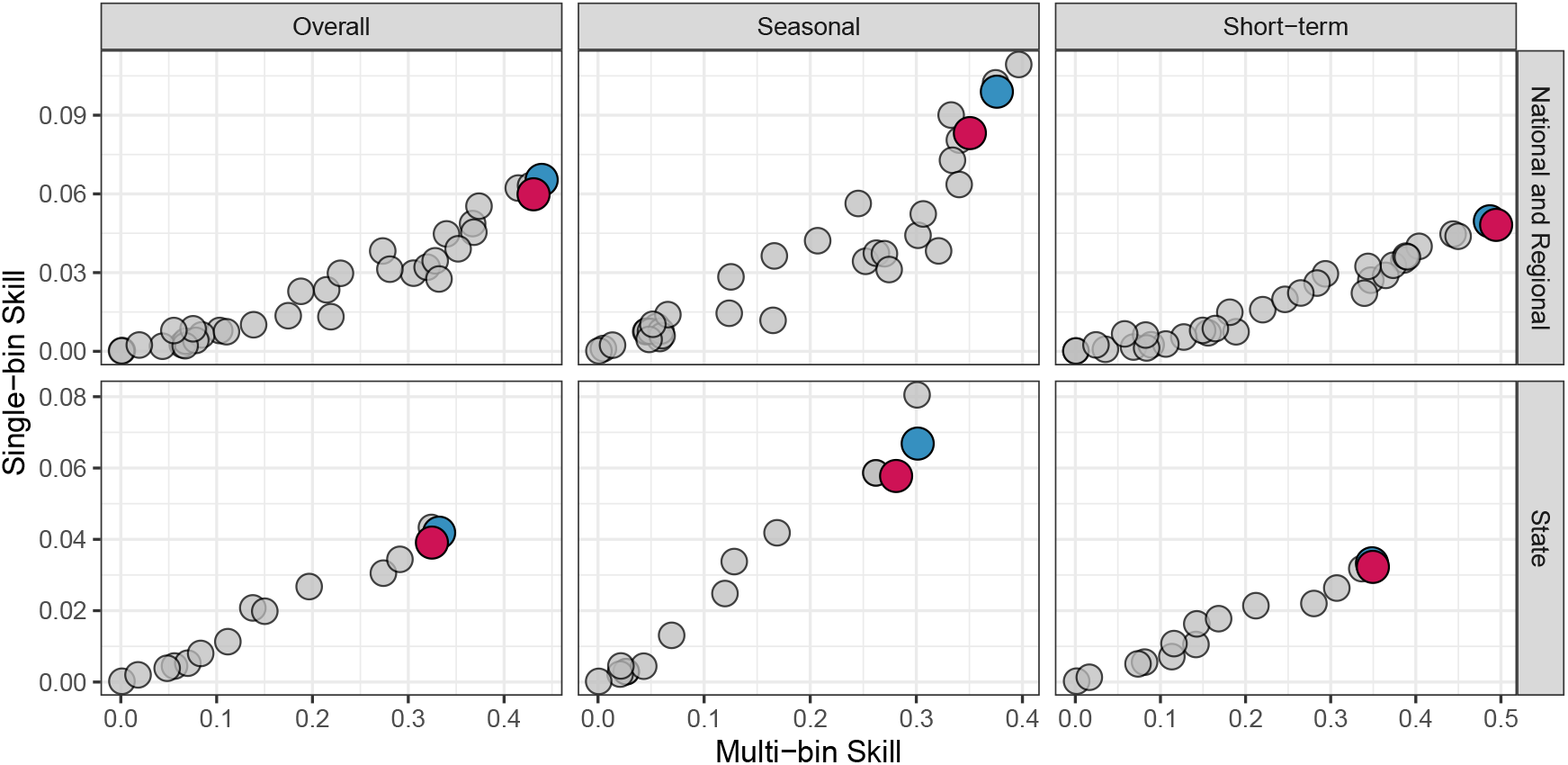
Results for the 2018/19 FluSight National and Regional challenge (top row) and State challenge (bottom row) for Inferno (red point), Dante (blue point) and all other models that participated in the 2018/19 FluSight challenges (grey points). The 2018/19 FluSight challenge evaluated models using multi-bin skill (x-axis), but starting with the FluSight 2019/20 challenge, will be using single-bin skill (y-axis). Skill scores are presented overall (left column), but also by seasonal targets (middle column) and short-term targets (right column). Inferno is a leading forecasting model overall, excelling in short-term forecasting, with good but not leading seasonal forecasting performance.

**Table 1:** The rank by challenge and target for Inferno and Dante as measured by single-bin and multi-bin skill. Inferno would have placed 2nd in both the National and Regional and the State challenges as measured by multi-bin skill, only finishing behind Dante. Inferno would have placed 4th (National and Regional) and 3rd (State) were the forecasts evaluated with single-bin skill. For both challenges and both evaluation metrics, Inferno achieved better short-term than seasonal performance.

| 2018/19 FluSight<br>Challenge | Target | Multi-bin Rank |  | Single-bin Rank |  |
| --- | --- | --- | --- | --- | --- |
|  |  | Inferno | Dante | Inferno | Dante |
| National and Regional | Overall | 2 | 1 | 4 | 1 |
|  | 1 wk ahead | 1 | 2 | 1 | 2 |
|  | 2 wk ahead | 1 | 2 | 2 | 1 |
|  | 3 wk ahead | 1 | 2 | 2 | 1 |
|  | 4 wk ahead | 2 | 1 | 2 | 1 |
|  | Season peak percentage | 5 | 1 | 5 | 3 |
|  | Season peak week | 11 | 8 | 11 | 8 |
|  | Season onset | 5 | 1 | 7 | 1 |
| State | Overall | 2 | 1 | 3 | 2 |
|  | 1 wk ahead | 3 | 1 | 3 | 1 |
|  | 2 wk ahead | 2 | 1 | 2 | 1 |
|  | 3 wk ahead | 1 | 2 | 2 | 1 |
|  | 4 wk ahead | 1 | 2 | 1 | 2 |
|  | Season peak percentage | 3 | 2 | 5 | 2 |
|  | Season peak week | 3 | 1 | 3 | 1 |

The small drop in predictive performance from Dante to Inferno is offset by Inferno’s significant improvement in runtime and preparation for future scalability to more granular forecasting geographies. Figure 11 shows the runtime comparison between Dante and Inferno at different stages of the flu season. Dante takes between 90 and 105 minutes to produce 25,000 MCMC samples throughout the 2018/19 flu season. Inferno takes between 30 seconds and 2 minutes to produce the same number of MCMC samples. The runtimes in Figure 11 are not directly comparable, as Inferno’s runtimes are for only one of the 64 geographies (53 states, 10 HHS regions, and the United States), while Dante’s runtimes are for all 64 geographies. If Inferno ran sequentially over all geographies, it would take roughly 30 minutes at the beginning of the season and 130 minutes at the end of the season, resulting in no computational gains over Dante by the end of the flu season. However, Inferno is trivially parallelizable. With cluster computing, all 64 geographies of Inferno could be computed simultaneously in two minutes or less. The significant advantage Inferno has over Dante is its scalability via parallelization. Thus, while Inferno’s predictive performance is comparable to but slightly worse than Dante’s, its significantly improved runtime and scalability make it a more attractive alternative for both the present and the future.

**Figure 11:**
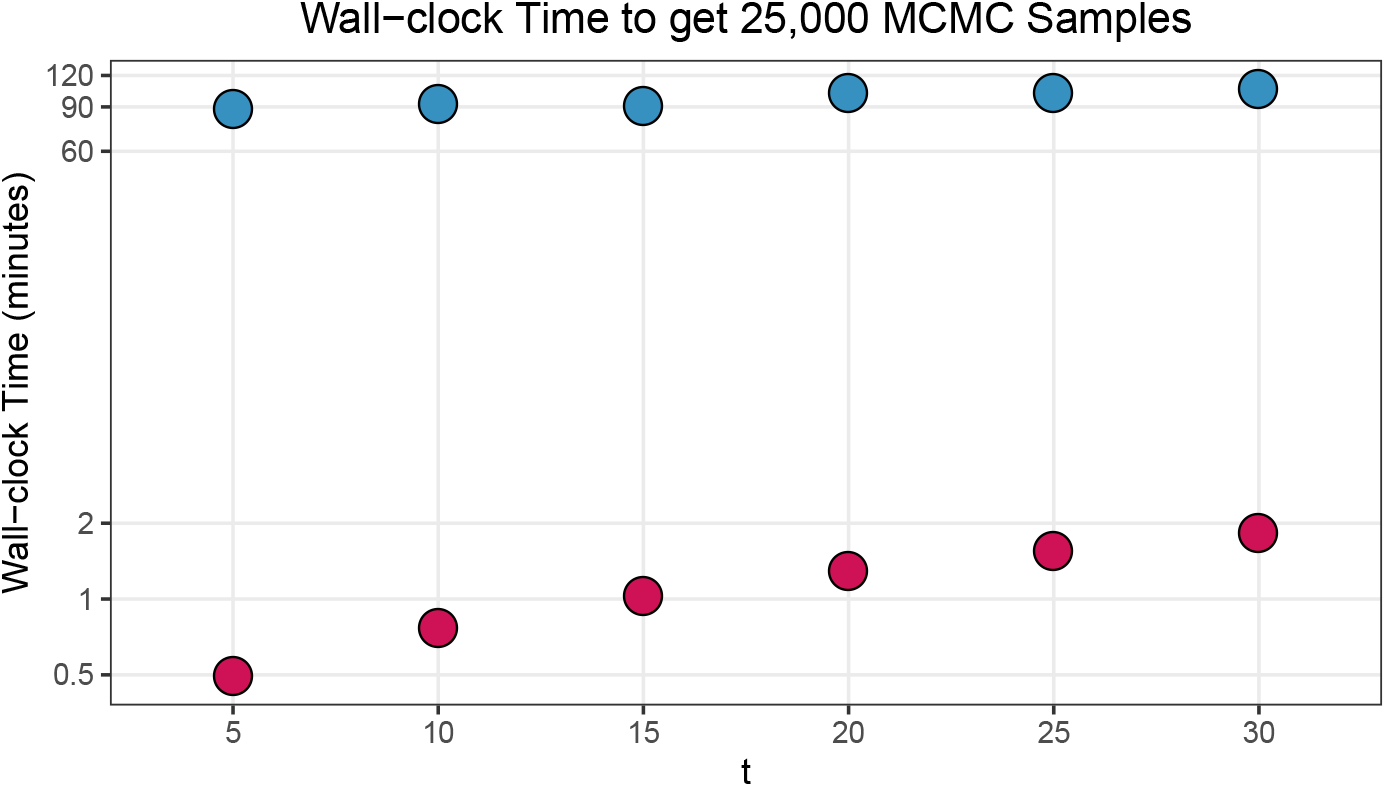
The average wall-clock runtime for a geography (e.g., a state) of Inferno (red) and actual wall-clock runtime for Dante (blue) to get 25,000 MCMC samples. Runtime increases as the size of the conditioning data increases for both Dante and Inferno. Inferno runs in 30 seconds early in the season (*t*=5) and takes approximately 2 minutes by the end of the flu season (*t*=30). In contrast, Dante takes approximately 90 minutes to run at the beginning of the flu season and 105 minutes by the end as a result of it conditioning on a much larger set of data.

## 4 Discussion

In this paper, I argued that while predictive performance is the most important measure of a forecasting model, it is not singularly important. Other factors like interpretability, generalizability, scalability, and runtime are also important. Developing a model with leading predictive performance but drastically improved runtime was the motivation behind Inferno. I laid out a six step procedure to estimate the parameters of Inferno from historical ILI data, greatly reducing the MCMC computations as executed by the probabilistic programming language JAGS. Furthermore, by forecasting each geography separately, Inferno can take advantage of parallelization, both improving forecast runtimes in the present while being scalable and well-positioned for the more spatially granular future of flu forecasting (e.g., county-level forecasting).

Inferno’s predictive performance was comparable to but worse than Dante’s. This may be for a couple different reasons, both of which are addressable. Firstly, Dante explicitly models backfill; previous work has shown that accounting for and modeling backfill can result in improved predictive performance [4, 11]. Similar modeling can be incorporated into Inferno at little additional computational cost. Secondly, Dante achieves self-consistency in its forecasts by modeling and forecasting all nested geographical units jointly. This self-consistency comes at a computational cost. The price Inferno pays to achieve significant computational speed-ups is the loss of self-consistency. There has been some recent work that takes independently generated probabilistic forecasts and, using principles of coherence, produces self-consistent forecasts that have improved predictive performance [8]. The combination of backfill modeling and coherence exploitation may result in equal or even better predictive performance at minimal computational cost.

Inferno’s development was motivated by the desire to build a forecasting model that maintains world-leading predictive performance while improving forecasting model properties not directly evaluated by FluSight. It is relatively straightforward to list characteristics we desire in an infectious disease forecasting model that extend well beyond predictive performance. For instance, we want forecasting models to

- reliably and accurately forecast public health relevant targets with actionable lead times
- quantify their forecast uncertainties
- incorporate public health interventions, facilitating “what-if” scenario assessments
- be transferrable between disease contexts and geographies
- run at all spatial scales
- be nimble and adaptable to ever-changing forecast settings
- run quickly, facilitating fast development and testing.

No forecasting model in existence today does all of these things well, but developing such a model should be the goal.

Forecasting challenges have proven to be highly effective organizing tools to help focus forecasters around a common goal, providing real value to public health responses. The singular goal of forecasters to maximize their predictive score is both a blessing and a curse. The predictive score helps generate competition which drives innovation and improvement. It also puts up blinders to all other characteristics we want forecasting models to have. A big question the forecasting community must address going forward is how forecasting challenges can be modified to expand their definition of what a “good” forecasting model is. That is, should forecasting challenges explicitly incorporate aspects of forecasting model generalizability, interpretability, utility, scalability, and speed and if so, how? Or are forecasting challenges not meant to assess forecasting models holistically but rather only assess one specific aspect of those models — their predictive performance?

## Acknowledgements

The author is thankful to C.C. Essix for her support and encouragement. This work was developed with the support of LANL’s LDRD research Project 20190546ECR. The author is thankful to the U.S. CDC FluSight team for making historical forecast submissions publicly available. Approved for unlimited release under LA-UR-20-30384.

## Appendix A

The following inputs are supplied to the JAGS model that implements Inferno:

- `T` is 35, the number of weeks of the flu season.
- `y` is a 35 × 1 vector where `y[t]` is the observed (w)ILI value for week `t` if it has been observed or `NA` if it has not. If `y[t]` is less than 0.0005 or greater than 0.9995, `y[t]` is set equal to 0.0005 or 0.9995, respectively.
- `alpha` is 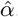, computed from Equation 14.
- `gamma` is a 35 × 1 vector where `gamma[t]` is 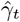, computed from Equation 19.
- `invCholUpper` is the inverse of the upper triangular Cholesky decomposition of 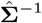, where 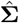 is computed from Equation 32 and 33.
- `sigma_mu` is the square root of 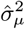, computed from Equation 24.

The JAGS code implementing Inferno is as follows:

~~~
model{
 for(t in 1:T){
 ## draw from posterior predictive distribution
 ypred[t] ∼ dbeta(alpha*theta[t], alpha*(1-theta[t]))
 ## data model
 y[t] ∼ dbeta(alpha*theta[t], alpha*(1-theta[t]))
 ## compute theta
 theta[t] <-ilogit(gamma[t] + delta[t])
}
 ## discrepancy GP model
 delta[1:T] <-mu + invCholUpper %*% Z[1:T]
 ## sample standard normals
 for(t in 1:T){
 Z[t] ∼ dnorm(0,1)
}
 ## discrepancy mean model
 mu ∼ dnorm(0,pow(sigma_mu,-2))
}
~~~

